# Statistical calibration of microbial suspensions in carrier controls during a textile disinfection ring trial

**DOI:** 10.1101/2020.11.19.389635

**Authors:** Antonio Monleon-Getino, AC Marca Home Care

## Abstract

**Introduction:** The high number of uncontrollable variables in microbiological systems increases experimental complexity and reduces accuracy, potentially leading to data misinterpretation or uncorrectable errors. During an interlaboratory calibration analysis it was observed that the microbial logarithmic reduction (LR) caused by disinfectants depends not only on the type of disinfectant but also on the initial microbial load in the fabric carriers, which can produce a misinterpretation of the results. Fabric carriers are commonly used in standard tests such as EN16616 and ASTM2274.

**Objective:** A method based on statistical calibration is proposed using a regression line between N_0_ (initial microbial load in the carrier) and LR to eliminate the influence of one on the other.

**Results:** An example with *Candida albicans* is presented. Once the method was applied, the influence of N_0_ on LR was eliminated and the new LR values can be used for factorial experiments, for example, to check the efficacy of disinfectants or detergents without depending on the microbial load placed in the carrier.

## 1. Background

The high number of variables at play in a microbiological study arises from the inherent complexity of living systems. Each variable contributes a certain degree of error that may propagate linearly or non-linearly, depending on the system. The more variables involved, the more errors can be expected, which can modify the outcome of a given study.

The causes of variability in microbiology methods can be roughly divided into three groups: the test system (e.g. the microorganism and environmental conditions), the scientist(s) performing the study, and in the case of antimicrobial efficacy studies, the test substance (1). Although several sources of variability have been determined, others that are unknown to the researchers may have an equally substantial effect (1). In general, experimental results must be reproducible to be meaningful. As this variability is known to exist in biological systems and is impossible to avoid, strategies should be sought to reduce its impact on significance.

In the field of textile disinfection [3], there are standard laboratory practices (EN16616; ASTM2274) that use microbiological systems to test the efficacy of a disinfectant product under conditions simulating use (P2S2 tests). These kinds of standards commonly involve the determination of the LR using artificially contaminated fabric carriers (N_0_). The biomonitors are laundered in a washing machine or laboratory-scale devices taking into account parameters such as contact time, temperature, and type of detergent or disinfectant.

LR is the difference in the common logarithm of the microbial count per mL of the initial load in the carriers before (N_0_) and after laundering (Na). In interlaboratory calibration analyses, LR is one of the main variables to test the accuracy and variability of experimental methods. For this reason, it is important to be very precise with the amount of microorganisms inoculated in the carrier or develop statistical resources to correct the correlation between N_0_ and LR [3][4].

### Example of the significant relationship between N_0_ and LR for *Candida albicans* observed during an interlaboratory calibration

Figure 1 depicts a significant correlation between N_o_ (Y axis) and LR (X axis). Axis?

**Figure 1:**
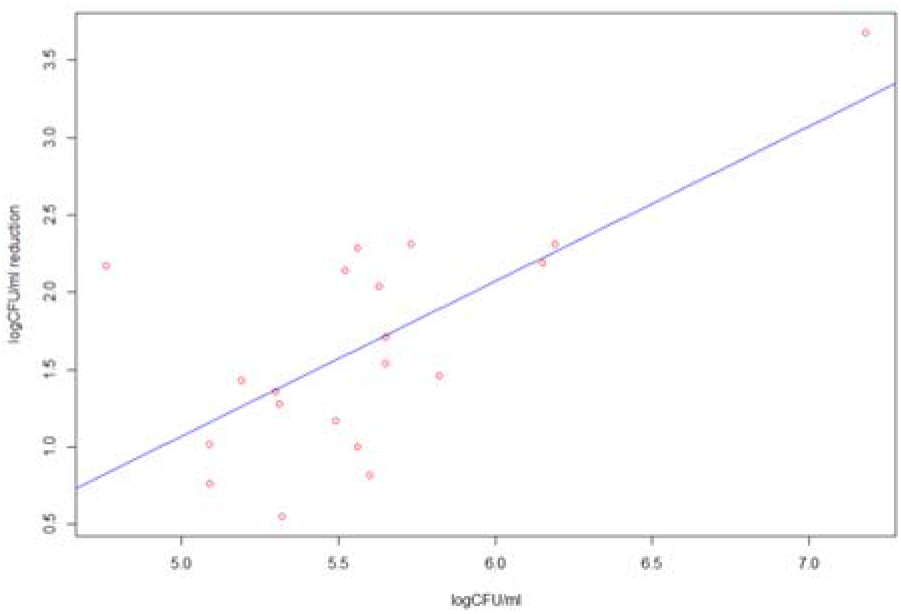
Regression between N_0_ (logCFU/ml) and reduction LR (logCFU/ml) in *C. albicans* was used.

### Results of the regression between N_0_ and LR

A regression was used to test a significant slope between N_0_ a nd LR, and in the results obtained using the lm() function of R [5] package it is possible to observe a significant correlation (p=0.000835) with a determination coefficient (R^2^=0.47):

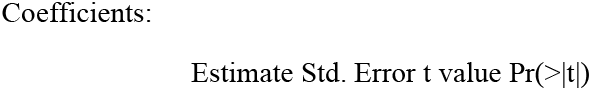
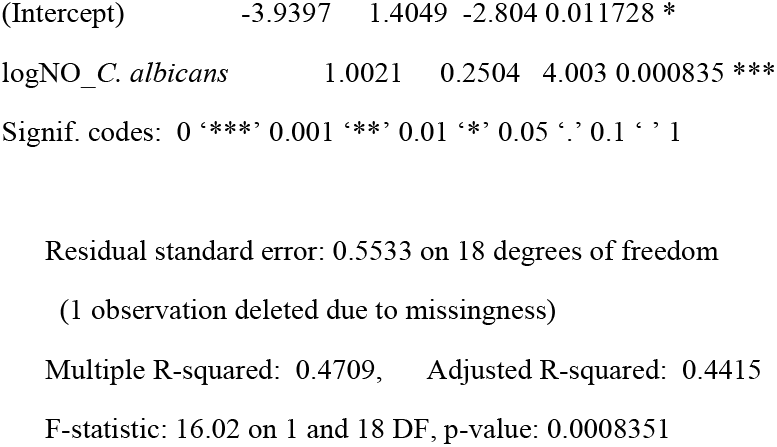

The regression used indicates a significant influence of the initial microbial load on the reduction of microorganisms in the carrier after laundering, which could lead to a misinterpretation of the LR value if not corrected.

## 2. Objectives

The general goal of this work was to develop a new methodology using regression and calibration that would avoid the influence between N_0_ and LR.

## 3. Material and methods

### Relationship (regression) Log CFU/ml reduction vs N_0_ using a statistical correction

Is the calibration line for N_0_ necessary? In this section, the regression methodology is used, in which a line is constructed between the LR values of the microorganisms and the value of their N_0_. For each microorganism, it is indicated if the model is significant and if so, it would suggest that a calibration is necessary. The scatterplot between the studied variables and the relationship line is also represented according to the literature [5][6].

We propose a correction [7][8][9] of the “reduction of log(CFU/ml)” variable in order to remove the influence of N_0_ on LR

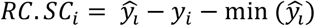

Where 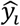 is the prediction of Log CFU/ml reduction based on a regression where the independent variable is 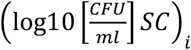 and a is the slope model and b is the independent term of the model: 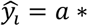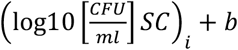

#### Relation between N_0_ and LR after calibration

After applying the proposed calibration, the regression line obtained is the one presented in Figure 2, where it can be seen that the slope is now 0 (p> 0.05, R^2^ approx = 0), providing no evidence of correlation.

**Figure 2:**
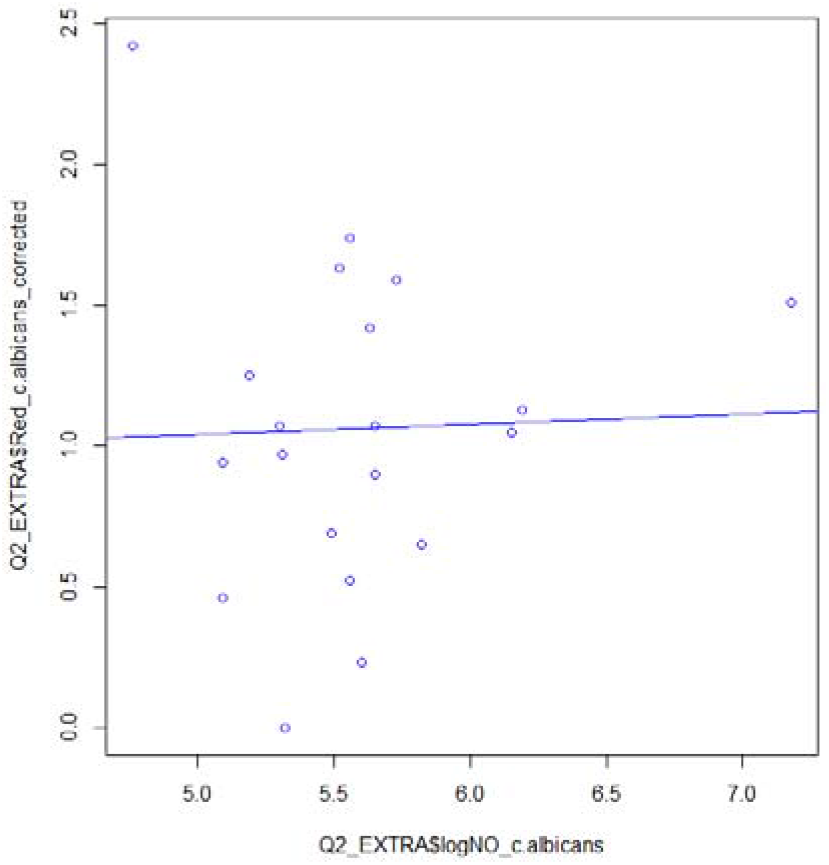
Regression between N_0_ (logCFU/ml) and reduction (logCFU/ml) in *C. albicans* is used.

#### CALIBRATION (CORRECTION OF REDUCTION IN FUNCTION OF N_0_)

##### Results of the regression (after calibration)

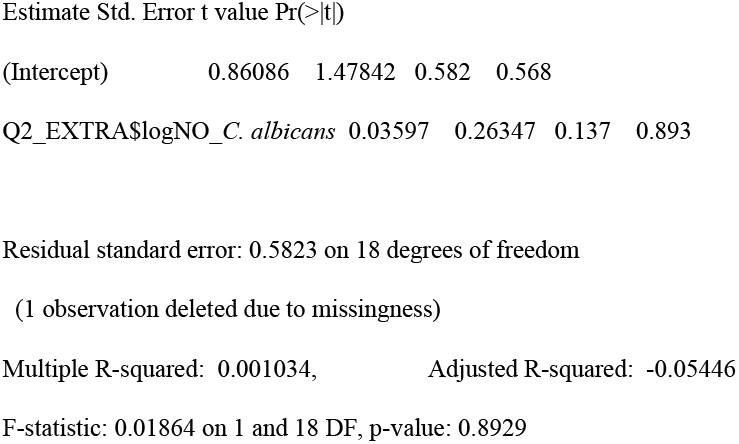

Once the influence of N_0_ on LR is removed, the new LR values can be used for factorial experiments, for example, to check the efficacy of disinfectants or sanitizers without depending on the dose of microorganism initially placed in the carrier.

## 5. Conclusions

In interlaboratory calibration analyses, LR is one of the main variables used to test the accuracy and variability of experimental methods. Microbial reduction caused by disinfectants depends not only on the type of disinfectant but also on the initial microbial load. A method based on statistical calibration using a regression line between N_0_ and LR is proposed to eliminate the influence of one on the other.

It was demonstrated using *C. albicans* as an example that once the calibration was applied, the influence of N_0_ on LR was eliminated. The new LR values can be used for factorial experiments, for example, to check the efficacy of disinfectants or detergents without depending on the microbial load initially placed in the carrier.

## References

[1] Microchem. 2020. Variability in Antimicrobial Testing (see at https://microchemlab.com/information/variability-antimicrobial-testing)

[2] Casadevall, A. and F.C. Fang, Reproducible science. Infect Immun, 2010. 78(12): p. 4972–5.

[3] Giacomini, A., et al., Experimental conditions may affect reproducibility of the beta-galactosidase assay. FEMS Microbiol Lett, 1992. 79(1–3): p. 87–90.

[4] McDonnell, G. and A.D. Russell, Antiseptics and disinfectants: activity, action, and resistance. Clin Microbiol Rev, 1999. 12(1): p. 147–79. 9. Ruano, M., J. El-Attrache, and P. Villegas, Efficacy comparisons of disinfectants used by the commercial poultry industry. Avian Dis, 2001. 45(4): p. 972–7.

[5] J Mendez, A Monleon-Getino, J Jofre, F Lucena. 2017. Use of non-linear mixed-effects modelling and regression analysis to predict the number of somatic coliphages by plaque enumeration after 3 hours of incubation. Journal of water and health 15 (5), 706–717.

[6] John Fox. 2010. Nonparametric Regression in R: An Appendix to An R Companion to Applied Regression, 2nd edition, revised December 2010“ (PDF). Socialsciences.mcmaster.ca.

[7] Loader, C. 2013 locfit: Local Regression, Likelihood and Density Estimation. R package version 1.5-9.1. http://CRAN.R-project.org/package=locfit

[8] Pinheiro, J., Bates, D., DebRoy, S., Sarkar, D. & R Core Team. 2015 nlme: Linear and Nonlinear Mixed Effects Models. R package version 3.1-120. http://CRAN.R-project.org/package=nlme

[9] T. Monleón-Getino. 2020. BDSbiost3: Machine learning and advanced statistical methods for omic, categorical analysis and others. Library for R published in github. Download in: https://github.com/amonleong/BDSbiost3

